# Alcohol Use Disrupts Age-Appropriate Cortical Thinning in Adolescence: A Data Driven Approach

**DOI:** 10.1101/2021.05.17.444458

**Authors:** Delin Sun, Viraj R. Adduru, Rachel D. Phillips, Heather C. Bouchard, Aristeidis Sotiras, Andrew M. Michael, Fiona C. Baker, Susan F. Tapert, Sandra A. Brown, Duncan B. Clark, David Goldston, Kate B. Nooner, Bonnie J. Nagel, Wesley K. Thompson, Michael De Bellis, Rajendra A. Morey

## Abstract

**Objective:** Cortical thickness changes dramatically during development and is influenced by adolescent drinking. However, previous findings have been inconsistent and limited by region-of-interest approaches that are underpowered because they do not conform to the underlying heterogeneity from the effects of alcohol.

**Methods:** Adolescents (n=657; 12-22 years at baseline) from the National Consortium on Alcohol and Neurodevelopment in Adolescence (NCANDA) who endorsed little to no alcohol use at baseline were assessed with structural MRI and followed longitudinally at four yearly intervals. Seven unique spatially covarying patterns of cortical thickness were obtained from the baseline scans by applying a novel data-driven method called non-negative matrix factorization (NMF). The cortical thickness maps of all participants’ longitudinal scans were projected onto vertex-level cortical patterns to obtain participant-specific coefficients for each pattern. Linear mixed-effects models were fit to each pattern to investigate longitudinal effects of alcohol consumption on cortical thickness.

**Results:** In most NMF-derived cortical thickness patterns, the longitudinal rate of decline in no/low drinkers was similar for all age cohorts, among moderate drinkers the decline was faster in the younger cohort and slower in the older cohort, among heavy drinkers the decline was fastest in the younger cohort and slowest in the older cohort (FDR corrected p-values < 0.01).

**Conclusions:** The NMF method can delineate spatially coordinated patterns of cortical thickness at the vertex level that are unconstrained by anatomical features. Age-appropriate cortical thinning is more rapid in younger adolescent drinkers and slower in older adolescent drinkers.

---

Neuromaturation occurs throughout childhood and adolescence while the brain undergoes dramatic changes in cortical gray-matter thickness and volume. Gray matter volume typically peaks before the teen years and then declines into adulthood as underutilized or redundant connections between neurons are pruned (1–3). Heavy adolescent alcohol use (4) is associated with faster cortical grey matter decline, possibly related to vulnerability during adolescent development (5). However, slower grey matter thinning has also been hypothesized on the premise that alcohol disrupts the process of pruning, which is typical in adolescence (4).

Alcohol is the most commonly misused substance with 24% of adolescents reporting consumption by 8^th^ grade and 59% before the end of high school (6). Most alarming is the number of drinks consumed by adolescents (7). While adolescents tend to drink less frequently and consume less overall than adults, they are much more likely to binge drink (8). The prevalence of binge drinking increases noticeably between 12-25 years (7) with 14% of 12^th^ graders binging once in two weeks (6).

Widespread differences in brain morphometry of adolescent drinkers have been identified (5, 9). Hypothesis-driven region of interest (ROIs) analyses have identified lower cortical thickness in the frontal, temporal, parietal, occipital, and cingulate cortices (2, 10, 11) of adolescent binge drinkers compared to light or non-drinking peers (12), an effect that was further amplified among younger adolescents (13, 14). However, competing findings of higher cortical thickness in the frontal, parietal, temporal, and occipital regions (13, 15) have also been reported. Previous studies have tested mean cortical thickness in ROIs based on anatomical definitions. However, the effects of binge drinking and patterns of brain maturation, which may interact with binge drinking, do not necessarily follow anatomical boundaries along prescribed gyral and sulcal features.

We addressed these challenges with a data-driven method called nonnegative matrix factorization (NMF) designed to identify patterns in cortical thickness variation at the vertex level, which is unconstrained by established anatomical borders (16). NMF is a class of multivariate algorithms that achieves matrix decomposition of an *m* x *n* input matrix (*m*: number of vertices; *n*: number of subjects) into two matrices with non-negative elements. Conceptually, NMF is similar to the more widely known method of independent component analysis (ICA).

We analyzed NCANDA data to assess the effects of adolescent binge drinking and its interactions with multiple risk factors, particularly age (17). NCANDA sampled a range of adolescent developmental periods within a shorter timeframe by adopting an *accelerated longitudinal study design*. Subjects from 12-22 years were recruited at baseline and followed longitudinally at yearly intervals for 5 years. We used NMF in the first stage to delineate covarying patterns of cortical thickness derived from baseline scans. In the second stage we used regression modeling to test alcohol-induced departures from normal developmental trajectories for each cortical thickness pattern. Consistent with prior evidence, we hypothesized that heavy drinking would be associated with more rapid age-related declines in cortical thickness (4).

## METHODS

### Participants

Adolescents (n=837; 12~22 years at baseline) were recruited from five sites: University of California at San Diego (*n*=214), Duke University (*n*=176), SRI International (*n*=169), Oregon Health and Science University (*n*=152), and University of Pittsburgh (*n*=126). Exclusionary criteria included serious medical, mental health, or learning disorders (17). Only youth who did not exceed drinking thresholds for alcohol (drinking class=0, see below; *n*=657) at baseline were enrolled (Fig. 1A) (17). Of the 657 participants assessed at baseline, 576 returned in 1 year for follow-up-1, 536 for follow-up-2, and 484 for follow-up-3, totaling 2,628 study visits.

**Figure 1.**
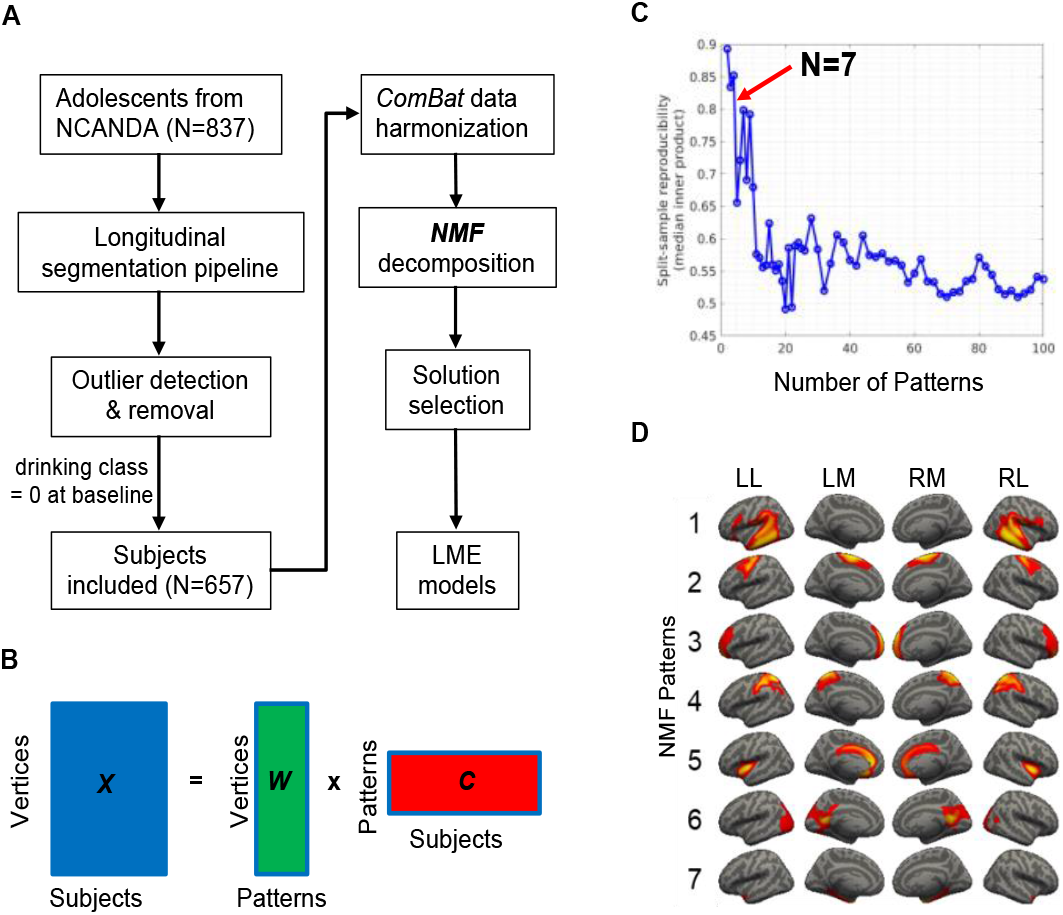
Analyses pipeline and non-negative matrix factorization (NMF) solutions. (A) Analyses pipeline. (B) NMF aims to find the solution that minimizes the difference between the raw data **X** and the reconstructed sample represented by the product of **W** and **C**. In matrix **X**, each row corresponds to a cortical vertex and each column corresponds to a subject. In matrix **W**, each row corresponds to the cortical thickness value of a vertex and each column corresponds to an NMF pattern. In matrix **C**, each row corresponds to a NMF pattern and each column corresponds to a subject. (C) The optimal number of NMF patterns is 7 based on peaks of the split-sample reproducibility analyses results (D) The optimal solution of 7 NMF patterns. Pattern 1 contains voxels in angular gyrus, supramarginal gyrus, inferior frontal areas, and superior/middle/inferior temporal regions; pattern 2 in superior and middle frontal regions; pattern 3 in frontopolar regions; pattern 4 in postcentral regions and superior parietal lobule; pattern 5 in anterior/middle cingulate cortex and bilateral insula; pattern 6 in posterior cingulate areas, lingual gyrus, cuneus, calcarine sulcus, and primary visual cortex; and pattern 7 in parahippocampal gyrus. LL = left hemisphere lateral view; LM = left hemisphere medial view; RM = right hemisphere medial view; RL = right hemisphere lateral view.

### Clinical and Demographic Measures

<u>Drinking class</u> reflects drinking behaviors at baseline and yearly thereafter measured with the Customary Drinking and Drug Use Record (CDDR) (17, 18) in four drinking classes: 1) no/low drinkers, 2) moderate drinker, 3) heavy drinkers, and 4) heavy drinkers. Detailed criteria for each drinking class are in Phillips et al (19).

<u>Self-identified race/ethnicity</u> included three categories: African-American, Caucasian, and Other.

<u>Socioeconomic status (SES)</u> was quantified using the highest years of education (range 6-20) of either parent (17) into low SES (6-12 years, n=47) and high SES (13-20 years, n=610).

<u>Family history of alcohol use and dependence (AUD) density</u> (range 0-4) was based on AUD in first- and second-degree relatives using the Family History Assessment Module (20).

<u>Cumulative trauma</u> was quantified as the sum of reported DSM-IV or DSM-5 Criterion-A traumatic events. DSM Criterion-A traumas and posttraumatic stress disorder (PTSD) symptoms as assessed separately in the subject and one parent at baseline as described in Phillips et al (19).

## MRI Acquisition and Longitudinal MRI Parcellation

All NCANDA sites collected MRI data on 3T GE MR750 or Siemens Tim Trio scanner using acquisition protocols for high-resolution structural imaging with 1-mm isotropic voxels as described in Phillips et al (19).

## Longitudinal Segmentation pipeline

Structural scans were processed using the FreeSurfer v6.0 longitudinal stream and longitudinal segmentation categorized into four steps: (1) cross-sectionally process, (2) create an unbiased template for each subject, (3) longitudinally process all timepoints, and (4) segment cortical regions. The longitudinal processing stream is described in Phillips et al (19). Spherical registration was performed to normalize the cortical thickness maps of 20,000 vertices in each subject to an average template. One participant was removed following failure of FreeSurfer longitudinal processing.

## Outlier Detection and Removal

Vertices whose cortical thickness was more than 3 times the standard deviation from the mean of all scans (all timepoints) were removed as outliers. A specific vertex could be excluded but the remaining vertices for the same subject at other timepoints were retained.

## Data Harmonization Across Sites

Harmonization of data from multiple sites/scanners was achieved by *ComBat*, a tool that models expected imaging features as linear combinations of the biological variables and scanner/site effects whose error term is further modulated by site-specific scaling factors (21). *ComBat* applies empirical Bayes to improve the estimation of site parameters by effectively removing unwanted sources of scanner/site variability while simultaneously increasing the power and reproducibility of subsequent statistical analyses of multi-site cortical thickness studies (21). The *ComBat* tool was used to harmonize cortical thickness values by removing scanner/site effects while preserving inherent biological associations such as age, sex, drinking class etc.

## NMF Decomposition

The spatial patterns in cortical thickness were estimated using NMF (16), which factors the data by positively weighting cortical thickness values that covary within the template and yields a highly specific and reproducible pattern-based representation. The goal of NMF is to find patterns of structural covariance that are common to all participants, such that a combination of these patterns with non-negative values approximates the original data. To achieve this goal, we first organized the cortical thickness data into a non-negative *m* x *n* matrix ***X*** with *m* vertices and *n* participants. We then represented the membership of the vertex-wise cortical thickness values to the patterns of structural covariance by a *m* x *v* matrix ***W*** where each row corresponds to the cortical thickness of a vertex and each column corresponds to a pattern. We also represented the contribution of each pattern to the whole cortical thickness map per participant with a *v* x *n* matrix ***C*** where each row corresponds to a pattern and each column corresponds to a participant. The NMF algorithm minimized the difference between the raw data ***X*** and the reconstructed sample represented by the product of ***W*** and ***C*** (Fig. 1B). Since matrix decomposition is generally not exactly solvable, we approximate it numerically with NMF. The cortical thickness values in the data matrix ***X*** that tend to covary are positively weighted, thus minimizing the reconstruction error and aggregating variance. The non-negativity constraint results in a non-overlapping pattern-based representation of whole-brain cortical thickness, which boasts advantageous specificity (16).

NMF was applied to cortical thickness maps from the baseline scans of 657 no/low drinkers to obtain basis vectors, which represent covariance patterns of cortical thickness to capture normal growth in adolescents with minimal drinking problems at baseline. The cortical thickness maps of all participants’ longitudinal scans were projected onto the basis vectors to obtain the participant-wise coefficients for each basis vector. NMF analysis was conducted with MATLAB scripts.

## Solution Selection

The NMF algorithm provides many possible solutions to the input matrix decomposition. Each solution contains a different number of covarying patterns. A solution with too many patterns may overfit the data by modeling noise fluctuations, while too few patterns may combine distinct patterns and thus undermine the model from expressing the underlying heterogeneity. We determined the appropriate number of patterns in the range of 2 to 100 based on reproducibility when the data was split into two sex- and age-matched halves (at baseline scan) (16). We calculated reproducibility across patterns by measuring the overlap between the independently estimated patterns from the inner product for the two splits using a combinatorial optimization algorithm known as the Hungarian Algorithm (22).

## Study Design of NCANDA

The goal of NCANDA was longitudinal study of age-related developmental brain changes associated with adolescent alcohol use. An *accelerated longitudinal design* was selected because it enabled studying our age range of interest in shorter time than a *single cohort design* (23) while investigating both within-subject and within-cohort structural brain changes. *Within-person age change* represents the difference between a subject’s age at each scan and their mean age across individual timepoints. Positive within-person age changes correspond to later visits while negative within-person age changes correspond to earlier visits by the same subject. *Cohort-age* represents the difference between a subject’s mean age across visits and the mean age of the entire sample across timepoints, thus centering cohort-age at the sample mean. Positive and negative cohort-ages refer to older and younger subjects respectively as compared to the sample mean. Each participant’s cohort-age remained constant across timepoints.

## Statistical Analyses

Following previously implemented approaches for structured multi-cohort longitudinal designs (23), we modeled the developmental trajectories of cortical thickness using a linear mixed-effects (LME) approach. In all models, participant identity was included as a random intercept to account for within-subject covariance across time. Drinking class, within-person age change, cohort-age, sex, self-identified ethnicity, SES, family history of AUD density, and cumulative lifetime trauma were included as fixed effects variables. We investigated the main effect of drinking class; the two-way interactions between drinking class and each of the other fixed effects variables; and three-way interactions between drinking class, within-person age change and cohort-age (without two-way interactions), on each of the NMF-derived patterns. The dependent variable for each regression model was the mean cortical thickness for all vertices contained in a given pattern. We also tested the main effects of within-person age change and cohort-age as well as their two-way interaction.

All statistical analyses were conducted in R using *lme4* for fitting LME models. The *sjPlot* package was used to plot the significant main and interaction effects. The false discovery rate (FDR) method (24) with a *q-value* threshold of 5% was applied to correct for multiple comparisons corresponding to each NMF pattern.

## RESULTS

### Clinical and Behavioral Results

See **Table 1** for sample demographic and alcohol use characteristics stratified by study visit. See **Table S1** for demographic characteristics organized by site at the baseline visit.

**Table 1.** Demographics at baseline and follow-ups

| Variable | Baseline<br>(n=657) | Follow-up 1<br>(n=576) | Follow-up 2<br>(n=536) | Follow-up 3<br>(n=484) |
| --- | --- | --- | --- | --- |
| **Drinking Class** |  |  |  |  |
| Low/No | 657 | 482 | 383 | 299 |
| Moderate | 0 | 68 | 93 | 105 |
| Heavy | 0 | 26 | 60 | 80 |
| **Sex** |  |  |  |  |
| Females | 329 | 288 | 265 | 246 |
| Males | 328 | 288 | 271 | 238 |
| **Age at Scan (Years)** |  |  |  |  |
| Mean | 15.6 | 16.8 | 17.7 | 18.7 |
| SD | 2.3 | 2.3 | 2.3 | 2.3 |
| **Self-declared Ethnicity** |  |  |  |  |
| Caucasian | 483 | 428 | 401 | 359 |
| African American | 92 | 77 | 73 | 63 |
| Other | 82 | 71 | 62 | 62 |
| **Socioeconomic Status** |  |  |  |  |
| 6-12 years | 47 | 37 | 37 | 36 |
| 13-20 years | 610 | 539 | 499 | 448 |
| **Family History Alcohol Density** |  |  |  |  |
| Mean Score | 0.20 | 0.00 | 0.17 | 0.16 |
| SD | 0.46 | 0.05 | 0.40 | 0.35 |
| **Cumulative Traumatic Events** |  |  |  |  |
| Mean | 1.01 | 1.01 | 1.00 | 1.00 |
| SD | 1.04 | 1.05 | 1.03 | 1.05 |

### NMF Patterns

We found the most prominent peak in split-sample reproducibility for the solution with 7 NMF patterns (Fig. 1C). Results associated with a smaller 9-pattern peak are reported in the Supplementary Materials. The results of the two solutions are concordant.

Nearly all patterns were symmetric bilaterally. As shown in Fig. 1D, pattern-1 contains voxels in angular gyrus, supramarginal gyrus, inferior frontal areas, and superior/middle/inferior temporal regions; pattern-2 is related to superior and middle frontopolar frontal regions; pattern-3 is associated with regions; pattern-4 is associated with postcentral regions and superior parietal lobule; pattern-5 is mainly associated with anterior/middle cingulate cortex and bilateral insula; pattern-6 is associated to posterior cingulate areas, lingual gyrus, cuneus, calcarine sulcus, and primary visual cortex; and pattern-7 is related to parahippocampal gyrus.

### Main Effect of Cohort-Age

Older cohort-age was associated with lower cortical thickness in all 7 patterns (*β*-values= −0.826~−0.127, *t*-values= −12.050~−2.574, *q*-values < 0.01; Table 2, Fig. S1).

**Table 2.** Main effects of drinking class, within-person age change, and cohort-age per NMF pattern.

| Pattern | $\beta$ | SD | df | t | p | q |
| --- | --- | --- | --- | --- | --- | --- |
| *Main effect of drinking class* |  |  |  |  |  |  |
| 1 | -0.152 | 0.081 | 1699.0 | -1.873 | 0.061 | 0.143 |
| 2 | -0.215 | 0.078 | 1696.3 | -2.752 | 0.006 | 0.042 |
| 3 | -0.110 | 0.092 | 1715.3 | -1.194 | 0.233 | 0.326 |
| 4 | -0.180 | 0.090 | 1713.1 | -2.014 | 0.044 | 0.143 |
| 5 | 0.032 | 0.062 | 1711.0 | 0.520 | 0.603 | 0.704 |
| 6 | -0.086 | 0.070 | 1705.9 | -1.221 | 0.222 | 0.326 |
| 7 | 0.001 | 0.050 | 1670.6 | 0.023 | 0.982 | 0.982 |
| *Main effect of within-person age change* |  |  |  |  |  |  |
| 1 | -1.382 | 0.032 | 1617.0 | -42.812 | <0.001 | <0.001 |
| 2 | -0.766 | 0.031 | 1615.5 | -24.647 | <0.001 | <0.001 |
| 3 | -1.373 | 0.037 | 1620.6 | -37.328 | <0.001 | <0.001 |
| 4 | -1.096 | 0.036 | 1618.4 | -30.649 | <0.001 | <0.001 |
| 5 | -0.956 | 0.025 | 1617.8 | -38.644 | <0.001 | <0.001 |
| 6 | -0.963 | 0.028 | 1616.1 | -34.401 | <0.001 | <0.001 |
| 7 | -0.241 | 0.020 | 1611.1 | -12.166 | <0.001 | <0.001 |
| *Main effect of cohort-age* |  |  |  |  |  |  |
| 1 | -0.702 | 0.069 | 658.5 | -10.219 | <0.001 | <0.001 |
| 2 | -0.396 | 0.067 | 657.2 | -5.933 | <0.001 | <0.001 |
| 3 | -0.826 | 0.073 | 659.7 | -11.342 | <0.001 | <0.001 |
| 4 | -0.672 | 0.071 | 657.4 | -9.468 | <0.001 | <0.001 |
| 5 | -0.596 | 0.049 | 657.1 | -12.050 | <0.001 | <0.001 |
| 6 | -0.638 | 0.057 | 656.0 | -11.177 | <0.001 | <0.001 |
| 7 | -0.127 | 0.049 | 656.9 | -2.574 | 0.010 | 0.010 |

Note: the regression models is “ $y \sim$ cohort-age + within-person age change + drinking class + sex + ethnicity + SES + family history of AUD density + life trauma + (1|participant ID)”. y is the mean cortical thickness of a pattern. $\beta$ , fixed-effect regression coefficient; SD, standard deviation; df, degree of freedom; t, t value; p, uncorrected p value; q, FDR corrected p value.

### Main Effect of Within-Person Age Change

Older within-person age was associated with lower cortical thickness in all 7 patterns (*β*-values= − 1.382~−0.241, *t*-values= −42.812~−12.166, *q*-values < 0.001; Table 2, Fig. S2).

### Main Effect of Drinking Class

Higher drinking class, indicative of heavier drinking, was associated with lower cortical thickness in pattern-2 (*β*-value= −0.215, *t*-value= − 2.752, *q*-value=0.042; Table 2, Fig. S3) but not the other patterns (*β*-values= −0.180~0.001, *t*-values= − 2.014~0.023, *q*-values > 0.1).

### Interaction of Within-Person Age Change and Cohort-Age

Significant interaction between within-person age change and cohort-age was found in all patterns (*β*-values=0.046~0.167, *t*-values=3.757~13.340, *q*-values < 0.001) except for pattern-7 (*β*-value=0.012, *t*-value=1.534, *q*-value=0.125). As shown in Table 3, Fig. S4, older cohorts had a slower rate of within-person age-related cortical thickness decline as compared to younger cohorts.

**Table 3.** Interaction effects among drinking class, within-person age change, and cohort-age per NMF-derived pattern.

| Pattern | $\beta$ | SD | df | t | p | q |
| --- | --- | --- | --- | --- | --- | --- |
| *Drinking class x within-person age change interaction <sup>1</sup>* |  |  |  |  |  |  |
| 1 | 0.031 | 0.075 | 1615.1 | 0.416 | 0.678 | 0.790 |
| 2 | -0.033 | 0.073 | 1613.5 | -0.458 | 0.647 | 0.790 |
| 3 | 0.126 | 0.086 | 1618.6 | 1.475 | 0.140 | 0.327 |
| 4 | 0.096 | 0.083 | 1616.5 | 1.156 | 0.248 | 0.433 |
| 5 | -0.002 | 0.058 | 1615.8 | -0.033 | 0.973 | 0.973 |
| 6 | 0.162 | 0.065 | 1614.1 | 2.480 | 0.013 | 0.093 |
| 7 | -0.073 | 0.046 | 1609.2 | -1.592 | 0.112 | 0.327 |
| *Drinking class x cohort-age interaction <sup>2</sup>* |  |  |  |  |  |  |
| 1 | 0.237 | 0.036 | 1687.5 | 6.629 | <0.001 | <0.001 |
| 2 | 0.135 | 0.035 | 1686.5 | 3.887 | <0.001 | <0.001 |
| 3 | 0.238 | 0.041 | 1702.9 | 5.817 | <0.001 | <0.001 |
| 4 | 0.249 | 0.040 | 1700.3 | 6.271 | <0.001 | <0.001 |
| 5 | 0.134 | 0.028 | 1699.3 | 4.847 | <0.001 | <0.001 |
| 6 | 0.213 | 0.031 | 1693.4 | 6.867 | <0.001 | <0.001 |
| 7 | 0.031 | 0.022 | 1663.6 | 1.386 | 0.166 | 0.166 |
| **Within-person age change x cohort-age interaction <sup>3</sup>** |  |  |  |  |  |  |
| 1 | 0.161 | 0.012 | 1597.5 | 13.339 | <0.001 | <0.001 |
| 2 | 0.046 | 0.012 | 1596.8 | 3.757 | <0.001 | <0.001 |
| 3 | 0.167 | 0.014 | 1598.5 | 11.990 | <0.001 | <0.001 |
| 4 | 0.153 | 0.014 | 1596.5 | 11.233 | <0.001 | <0.001 |
| 5 | 0.106 | 0.009 | 1596.1 | 11.227 | <0.001 | <0.001 |
| 6 | 0.140 | 0.011 | 1595.1 | 13.340 | <0.001 | <0.001 |
| 7 | 0.012 | 0.008 | 1597.0 | 1.534 | 0.125 | 0.125 |
| **Drinking class x within-person age change x cohort-age interaction <sup>4</sup>** |  |  |  |  |  |  |
| 1 | 0.161 | 0.024 | 1621.7 | 6.810 | <0.001 | <0.001 |
| 2 | 0.065 | 0.023 | 1620.7 | 2.842 | <0.001 | <0.001 |
| 3 | 0.182 | 0.027 | 1626.5 | 6.786 | <0.001 | <0.001 |
| 4 | 0.166 | 0.026 | 1624.4 | 6.347 | <0.001 | <0.001 |
| 5 | 0.094 | 0.018 | 1623.9 | 5.152 | <0.001 | <0.001 |
| 6 | 0.163 | 0.020 | 1621.1 | 8.002 | <0.001 | <0.001 |
| 7 | 0.011 | 0.015 | 1614.6 | 0.725 | 0.469 | 0.469 |

Note: regression models are
1, $y \sim \text{cohort-age} + \text{within-person age change} * \text{drinking class} + \text{sex} + \text{ethnicity} + \text{SES} + \text{family history of AUD density} + \text{life trauma} + (1 | \text{participant ID})$ ;
2, $y \sim \text{within-person age change} + \text{cohort-age} * \text{drinking class} + \text{sex} + \text{ethnicity} + \text{SES} + \text{family history of AUD density} + \text{life trauma} + (1 | \text{participant ID})$ ;
3, $y \sim \text{within-person age change} * \text{cohort-age} + \text{drinking class} + \text{sex} + \text{ethnicity} + \text{SES} + \text{family history of AUD density} + \text{life trauma} + (1 | \text{participant ID})$ ;
4, $y \sim \text{within-person age change} : \text{cohort-age} : \text{drinking class} + \text{sex} + \text{ethnicity} + \text{SES} + \text{family history of AUD density} + \text{life trauma} + (1 | \text{participant ID})$ .
$y$ is the mean cortical thickness of a pattern. $\beta$ , fixed-effect regression coefficient; SD, standard deviation; df, degree of freedom; $t$ , $t$ value; $p$ , uncorrected $p$ value; $q$ , FDR corrected $p$ value.

### Interaction of Drinking Class and Cohort-Age

Significant interactions were found in all patterns (*β*-values=0.134~0.249, *t*-values=3.887~6.867, *q*-values < 0.001) except pattern-7 (*β*-value=0.031, *t*-value=1.386, *q*-value=0.166). As shown in Table 3, Fig. S5, higher drinking class was associated with a slower rate of cohort-age-related cortical thickness decline.

### Interaction of Drinking Class and Within-Person Age Change

No significant result was found in any pattern (*β*-values= −0.073~0.162, *t*-values= −1.592~2.480, *q*-values > 0.09; Table 3).

### Interaction of Drinking Class, Cohort-age and Within-Person Age Change

Significant three-way interactions were found in all patterns (*β*-values=0.065~0.182, *t*-values=2.842~8.002, *q*-values < 0.01) except for pattern-7 (*β*-value=0.011, *t*-value=0.725, *q*-value=0.469). As shown in Table 3 and Fig. 2, the longitudinal rate of cortical thickness decline in no/low drinkers was similar for all age cohorts, decline among moderate drinkers was faster in the younger cohort and slower in the older cohort, decline among heavy drinkers was fastest in the younger cohort and slowest in the older cohort.

**Figure 2.**
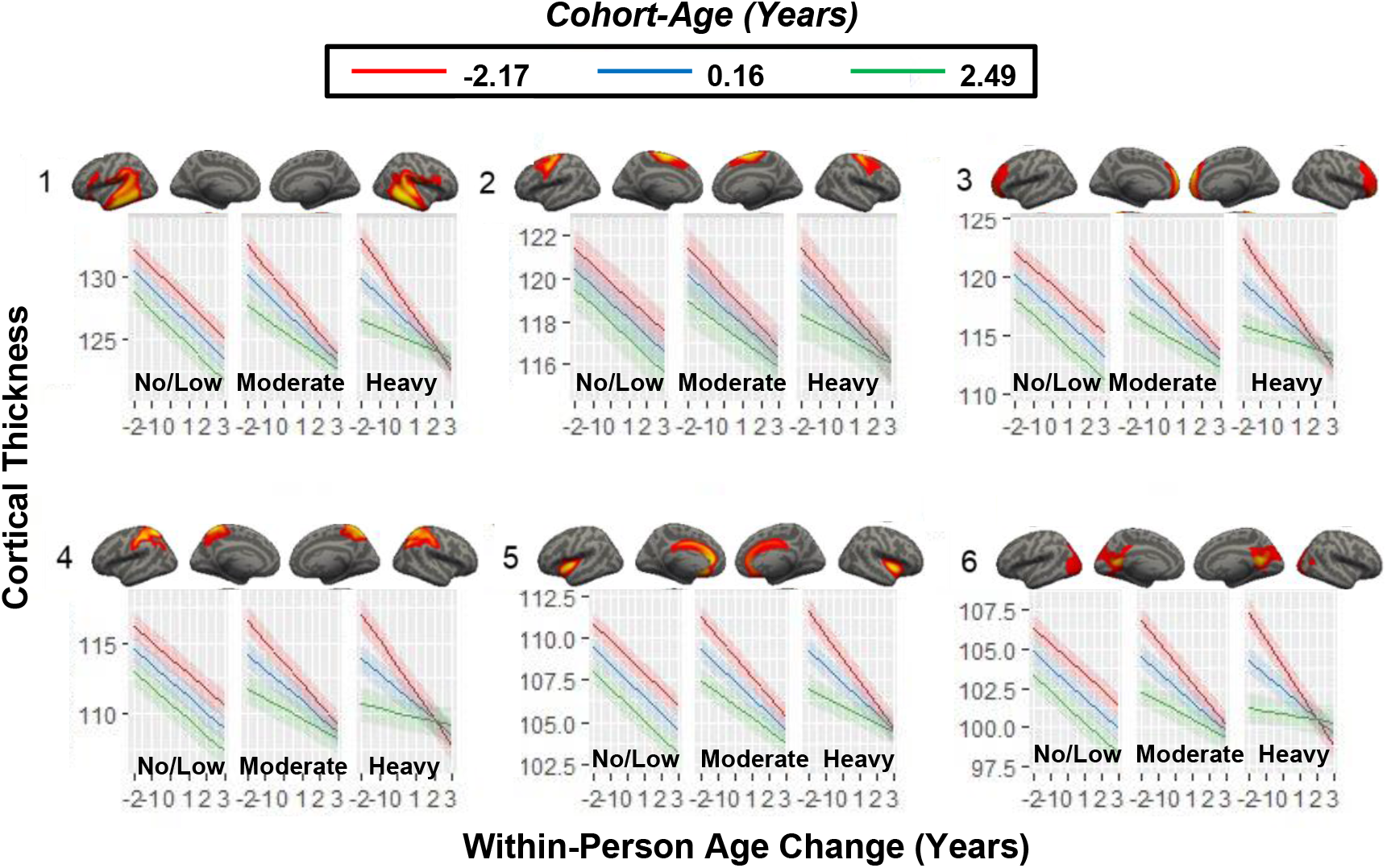
Three-way interactions between drinking class, cohort-age and within-person age change. The rates of within-person age-related cortical thickness declines are similar across age cohorts in no/low drinkers, faster from the younger cohort and slower from the older cohort in moderate drinkers, and fastest from the younger cohort and slowest from the older cohort in heavy drinkers in all patterns (*β*-values = 0.065~0.182, *t*-values = 2.842~8.002, *q*-values < 0.01) except for pattern 7 (*β*-value = 0.011, *t*-value = 0.724, *q*-value = 0.469).

### Interaction of Drinking Class and Other Variables

There was no significant interaction between drinking class and the other variables including sex, cumulative trauma, self-identified ethnicity, SES, family history of AUD.

## DISCUSSION

We investigated the longitudinal effects of alcohol use on cortical development in a large adolescent sample. We applied NMF, a multivariate data-driven method, to cluster vertices into 7 covarying patterns of cortical thickness. Significant decline in cortical thickness that was modulated by cohort-age and longitudinal progression (within-person age change) in 6 of 7 cortical thickness patterns. Participants in older cohorts, relative to younger cohorts, experienced slower declines in cortical thickness over longitudinal visits. Adolescents who engaged in heavier drinking exhibited lower cortical thickness in superior and middle frontal regions, and experienced slower cohort-age associated declines in cortical thickness across widely distributed brain areas including frontal, parietal, temporal and cingulate cortices. Of particular interest, the longitudinal rate of cortical thickness decline was similar for all age cohorts among no/low drinkers in widely distributed regions over study visits. However, moderate and heavy drinking differentially disrupted the trajectory of cortical thinning across age cohorts. Specifically, cortical thinning among moderate drinkers was faster in the younger cohort and slower in the older cohort, whereas cortical thinning among heavy drinkers was fastest in the younger cohort and slowest in the older cohort (Figure 2).

Our results confirms that heavy drinking speeds-up the cortical thinning process, which is typical of adolescent brain development, particularly in younger adolescents and attenuates this process in older individuals (14). The present study is the largest to date to investigate cortical gray matter changes related to adolescent alcohol use and the first study to deploy an advanced data-driven multivariate method that is capable of identifying coordinated patterns of cortical thickness that are unconstrainted by anatomical boundaries (16). A small number of patterns are more sensitive at capturing statistically significant results compared to region-of-interest methods, which are unable to provide spatially targeted results, and whole brain vertex-wise analysis (14), which must survive stringent corrections for multiple comparisons. Consequently, both methods suffer from Type II error. More importantly, the patterns we identified are closely related to meaningful functional networks that recapitulate established patterns in large normative adolescent samples (16). Pattern-1 (Fig. 1D) is aligned with the frontoparietal network (FPN; also known as central executive network, CEN) and parts of the temporal cortex. The FPN is known for coordinating goal-directed behaviors in a rapid, accurate, and flexible manner (25). Prior evidence supports that adolescent binge drinking is linked to reduced prefrontal activation during executive processing (26, 27), weaker frontoparietal connectivity (28), and impaired executive function, which is reliant on FPN (29). Pattern-5, which includes the anterior and middle cingulate cortices and bilateral insula, are part of the salience network (SN). The SN plays a crucial role in consciously integrating autonomic feedback and responses with internal goals and external demands (30). Pattern-5 comports with evidence that adolescent binge drinking is linked to impaired response inhibition, which is heavily reliant on SN (31, 32). Adolescents who engage in binge drinking require increased effort when inhibiting prepotent responses to alcohol-related stimuli by engaging bilateral anterior insula and inferior frontal gyrus, both of which are core components of the SN (33, 34), as well as intact FPN function. The posterior cingulate cortex and precuneus within pattern-6 and the anterior cingulate cortex within pattern-5 are key components of the default mode network (DMN). The DMN is involved in self-referential processing, theory of mind (ToM), memory, and learning (35), which is active in the absence of goal-directed activity. Notably, the DMN is dysregulated in adolescents with family history of AUD (36) and adolescent binge drinkers have heightened connectivity between DMN regions, which impedes the maturation of affective and self-reflective neural systems and undermines the development of complex social and emotional behaviors (37, 38). Thus, our findings in FPN, SN, and DMN, networks integral to adolescent brain development (39–41), are consistent with evidence that adolescent alcohol exposure alters network activity and reconfigures network connectivity (42, 43). Our NMF-derived findings may guide hypothesis-driven investigations of alcohol effects on adolescent brain development.

The longitudinal rate of cortical thickness decline was similar for all age cohorts in no/low drinkers. However, moderate and heavy drinking differentially disrupted this trajectory of cortical thinning across age cohorts. Specifically, cortical thinning was faster in the younger cohort and slower in the older cohort among moderate drinkers, and fastest in the younger cohort and slowest in the older cohort among heavy drinkers in 6 patterns (Table 3, Fig. 2). The typical pattern of (within-person) age-related cortical thinning, which is more rapid in younger cohorts than older cohorts, is more disparate in moderate drinkers, and most disparate in heavy drinkers. Greater frequency and intensity of alcohol use appears to effect older cohorts by inducing more powerful neurotoxic effects in early adolescence and/or more profoundly delaying brain maturation via pruning in late adolescence. By contrast, alcohol adversely effects younger cohorts by delaying cortical maturation in early adolescence and/or eliciting severe neurotoxic effects in late adolescence. Relatedly, Squeglia et al (4) reported faster grey matter reduction only in adolescents engaged in heavy drinking. Previously published studies using anatomically defined ROIs obscured the underlying heterogeneity in age-related cortical thickness changes that may be associated with synaptic pruning that were revealed by our data-driven approach. It is also conceivable that cortical areas with robust pruning during adolescence are particularly susceptible to disruption from heavy alcohol consumption (44).

We found no significant interactions between drinking class and the other variables including lifetime trauma, SES, ethnicity, sex, and family history of alcohol use disorder. In a related NCANDA study, baseline PTSD symptoms but not the number of baseline traumatic events predicted moderate to heavy drinking longitudinally (45). A previous NCANDA study also reported similar age-related declines in cortical thickness in male and female adolescents and similar cortical thickness across races, suggesting that cortical thickness decline is typical of development (2). We reported that cumulative lifetime trauma and alcohol use interact to affect the volume and trajectory of hippocampal and amygdala substructures (19). It is possible that specific variables in adolescent drinkers, such as sex and trauma have stronger effects on cortical volume, surface area, and subcortical volume than on cortical thickness. Studies in small samples reported thinner cortices in adolescent male binge drinkers (N=14) and thicker cortices in adolescent female binge drinkers (N=15) compared to sex-matched alcohol-naïve controls and thinner cortices in frontal and parietal lobes in adolescents with a family history of AUD (N=95) than controls, especially among the youngest adolescents Previously reported interactions between alcohol consumption and sex (13), or family history (46) in relatively small samples with specific clinical and demographic attributes may not generalize to our much larger and more representative sample. Large-scale studies of adolescent cortical development that assess the role of alcohol with other environmental insults such as childhood trauma, poverty, drug use, and education may prove to be informative.

### Strengths and Limitations

A limitation of our study was that we did not control for cannabis or other substance use, but these occurred at extremely low levels in our sample. In addition, we analyzed the first four years of NCANDA data, but another time point that became available as we concluded data analysis will expand the sample size within each developmental cohort and may help define longer-term sequelae. Finally, our analysis draws only on cortical thickness data, but did not examine cortical surface area or white matter. Future studies that apply multi-modal imaging may discover novel effects of alcohol in the adolescent brain to inform treatment development.

Our study has several strengths relative to previous investigations. First, we applied unsupervised machine learning to achieve clustering, data dimension reduction, and enhanced power. Second, we investigated three times more participants than previous studies. Third, we utilized *ComBat* to harmonize cortical thickness measurements across five NCANDA sites while preserving variance associated with biologically and behaviorally relevant variables (21). Finally, we leveraged longitudinal data with four yearly timepoints, which is rare among neuroimaging studies of psychiatric conditions.

## Conclusions

Multivariate data-driven method can successfully identify coordinated patterns of vertex-level cortical thickness variation. Age-related cortical thinning, which is typical of the adolescent neurodevelopment process, occurs more rapidly in younger individuals and less rapidly in older individuals who engage in heavy alcohol consumption as compared to low/non-drinking adolescents. The present findings in the cortex mean that early adolescent binge drinking may have related adverse effects on wide ranging processes of cognition, emotion regulation, impulsivity, and social learning.

## AUTHOR AND ARTICLE INFORMATION

Duke University School of Medicine (Sun, Adduru, Phillips, Bouchard, Michael, Goldston, De Bellis, Morey); Washington University Department of Radiology and Institute for Informatics (Sotiras); SRI International Biosciences Division (Baker); University of California San Diego Department of Psychiatry (Tapert, Brown); University of Pittsburgh Department of Psychiatry (Clark); University of North Carolina Wilmington (Nooner); Oregon Health Sciences University Department of Psychiatry and Behavioral Sciences (Nagel).

Supported by NIH Grants AA021697, AA021695, AA021692, AA021696, AA021681, AA021690, and AA021691, Department of Veterans Affairs Mid-Atlantic MIRECC.

Mary Nicole Buckley and Molly Monsour assisted in the visual inspection of segmentations.

Data used here are from the data release NCANDA_PUBLIC_3Y_REDCAP_V03 (https://doi.org/10.7303/syn23702728) and NCANDA_PUBLIC_3Y_STRUCTURAL_V01 (https://doi.org/10.7303/syn22213272) distributed to the public according to the NCANDA Data Distribution agreement (www.niaaa.nih.gov/research/major-initiatives/national-consortium-alcohol-and-neurodevelopment-adolescence/ncandadata).

