## Supplementary materials for "Alcohol Use Disrupts Age-Appropriate Cortical Thinning in Adolescence: A Data Driven Approach"

***Supplementary Table S1*. Demographic Characteristics at Baseline by Cohort.**

|  | ***Pittsburgh***  *(*n*=92)* | ***SRI***  *(*n*=126)* | ***Duke***  *(*n*=140)* | ***OHSU***  *(*n*=130)* | ***UCSD*** *(*n*=169)* |
| --- | --- | --- | --- | --- | --- |
| ***Sex***  *Females*  *Males* | 50  42 | 59  67 | 73  67 | 65  65 | 82  87 |
| ***Age at Scan (Years)***  *Mean*  *SD* | 16.2  2.5 | 15.0  1.9 | 15.2  1.9 | 16.1  2.6 | 15.8  2.4 |
| ***Self-declared Ethnicity***  *Caucasian*  *African American*  *Other* | 71  19  2 | 98  1  27 | 76  55  9 | 109  3  18 | 129  14  26 |
| ***Socioeconomic Status***  *6-12 years*  *13-20 years* | 7  85 | 4  122 | 12  128 | 7  123 | 17  152 |
| ***Family History Alcohol Use Density***  *Mean Score*  *SD* | 0.12  0.30 | 0.24  0.60 | 0.09  0.24 | 0.22  0.43 | 0.31  0.54 |
| ***Cumulative Traumatic Events****  *Mean*  *SD* | 0.87  1.13 | 0.52  0.80 | 1.41  0.94 | 1.20  1.21 | 0.99  0.98 |

*, No participant met DSM criteria for PTSD at baseline. Pittsburg = University of Pittsburgh Medical Center; SRI = SRI International; Duke = Duke University Medical Center; OHSU = Oregon Health and Science University; UCSD = University of California at San Diego.

***Supplementary Table S2*. Interaction effects between drinking class and the other variables per NMF pattern.**

| **Pattern** | ***β*** | **SD** | **df** | ***t*** | ***p*** | ***q*** |
| --- | --- | --- | --- | --- | --- | --- |
|  | ***Drinking class x life trauma interaction*** ^1^ | | | |  |  |
| 1 | -0.013 | 0.069 | 1683.3 | -0.193 | 0.847 | 0.847 |
| 2 | -0.039 | 0.067 | 1680.9 | -0.578 | 0.563 | 0.847 |
| 3 | 0.025 | 0.079 | 1697.4 | 0.315 | 0.752 | 0.847 |
| 4 | -0.027 | 0.077 | 1695.4 | -0.348 | 0.728 | 0.847 |
| 5 | 0.057 | 0.053 | 1693.5 | 1.070 | 0.285 | 0.847 |
| 6 | -0.030 | 0.060 | 1689.0 | -0.495 | 0.620 | 0.847 |
| 7 | -0.062 | 0.043 | 1658.9 | -1.444 | 0.149 | 0.847 |
|  | ***Drinking class x sex interaction*** ^2^ | | | |  |  |
| 1 | -0.114 | 0.146 | 1693.2 | -0.784 | 0.433 | 0.606 |
| 2 | -0.089 | 0.140 | 1690.6 | -0.632 | 0.528 | 0.616 |
| 3 | -0.027 | 0.166 | 1708.8 | -0.162 | 0.871 | 0.871 |
| 4 | -0.191 | 0.161 | 1706.6 | -1.187 | 0.235 | 0.536 |
| 5 | 0.114 | 0.111 | 1704.4 | 1.024 | 0.306 | 0.536 |
| 6 | -0.275 | 0.126 | 1699.6 | -2.185 | 0.029 | 0.203 |
| 7 | 0.101 | 0.090 | 1666.0 | 1.132 | 0.258 | 0.536 |
|  | ***Drinking class x ethnicity interaction*** ^3^ | | | |  |  |
| 1 | 0.075 | 0.277 | 1687.2 | 0.271 | 0.786 | 0.917 |
| 2 | -0.280 | 0.266 | 1684.3 | -1.050 | 0.294 | 0.686 |
| 3 | 0.099 | 0.315 | 1702.1 | 0.314 | 0.754 | 0.917 |
| 4 | -0.418 | 0.306 | 1699.6 | -1.366 | 0.172 | 0.686 |
| 5 | 0.008 | 0.212 | 1697.9 | 0.037 | 0.970 | 0.970 |
| 6 | -0.117 | 0.240 | 1693.2 | -0.489 | 0.625 | 0.917 |
| 7 | 0.210 | 0.170 | 1661.5 | 1.234 | 0.217 | 0.686 |
|  | ***Drinking class x SES interaction*** ^4^ | | | |  |  |
| 1 | -0.221 | 0.318 | 1687.0 | -0.694 | 0.488 | 0.792 |
| 2 | 0.129 | 0.306 | 1684.6 | 0.420 | 0.675 | 0.792 |
| 3 | -0.350 | 0.361 | 1701.7 | -0.968 | 0.333 | 0.792 |
| 4 | -0.048 | 0.352 | 1699.6 | -0.137 | 0.891 | 0.891 |
| 5 | -0.184 | 0.243 | 1697.7 | -0.757 | 0.449 | 0.792 |
| 6 | -0.114 | 0.276 | 1693.0 | -0.414 | 0.679 | 0.792 |
| 7 | -0.177 | 0.196 | 1661.6 | -0.903 | 0.367 | 0.792 |
|  | ***Drinking class x family history of AUD density interaction*** ^5^ | | | | |  |
| 1 | 0.202 | 0.177 | 1642.6 | 1.142 | 0.254 | 0.493 |
| 2 | 0.117 | 0.170 | 1640.8 | 0.689 | 0.491 | 0.493 |
| 3 | 0.228 | 0.201 | 1650.5 | 1.135 | 0.257 | 0.493 |
| 4 | 0.149 | 0.196 | 1648.4 | 0.762 | 0.446 | 0.493 |
| 5 | 0.129 | 0.135 | 1647.2 | 0.954 | 0.340 | 0.493 |
| 6 | 0.116 | 0.153 | 1644.4 | 0.760 | 0.448 | 0.493 |
| 7 | 0.074 | 0.108 | 1629.2 | 0.686 | 0.493 | 0.493 |

Note: regression models are

^1^, y ~ cohort-age + within-person age change + drinking class * life trauma + sex + ethnicity + SES + family history of AUD density + (1|participant ID);

^2^, y ~ cohort-age + within-person age change + drinking class * sex + life trauma + ethnicity + SES + family history of AUD density + (1|participant ID);

^3^, y ~ cohort-age + within-person age change + drinking class * ethnicity + life trauma + sex + SES + family history of AUD density + (1|participant ID);

^4^, y ~ cohort-age + within-person age change + drinking class * SES + life trauma + sex + ethnicity + family history of AUD density + (1|participant ID);

5, y ~ cohort-age + within-person age change + drinking class * family history of AUD density + life trauma + sex + ethnicity + SES + (1|participant ID).

y is the mean cortical thickness of a pattern. *β*, fixed-effect regression coefficient; SD, standard deviation; df, degree of freedom; *t*, t value; *p*, uncorrected *p* value; *q*, FDR corrected *p* value.

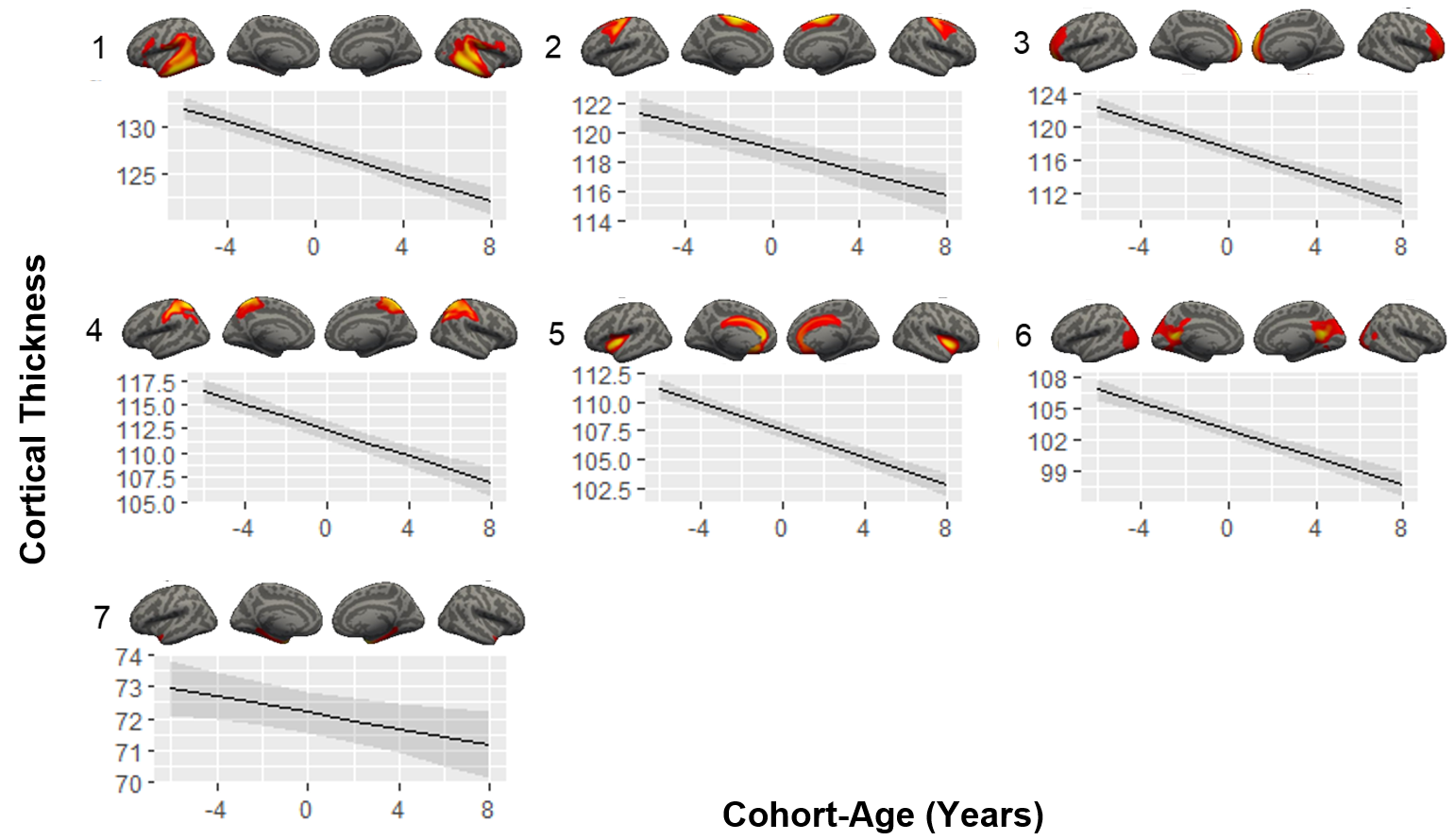
***Supplementary Figure S1.*** Main effect of cohort-age. Older cohort-age is associated with lower cortical thickness in all 7 patterns (*β*-values = -0.826~-0.127, *t*-values = -12.050~-2.574, *q*-values < 0.010).

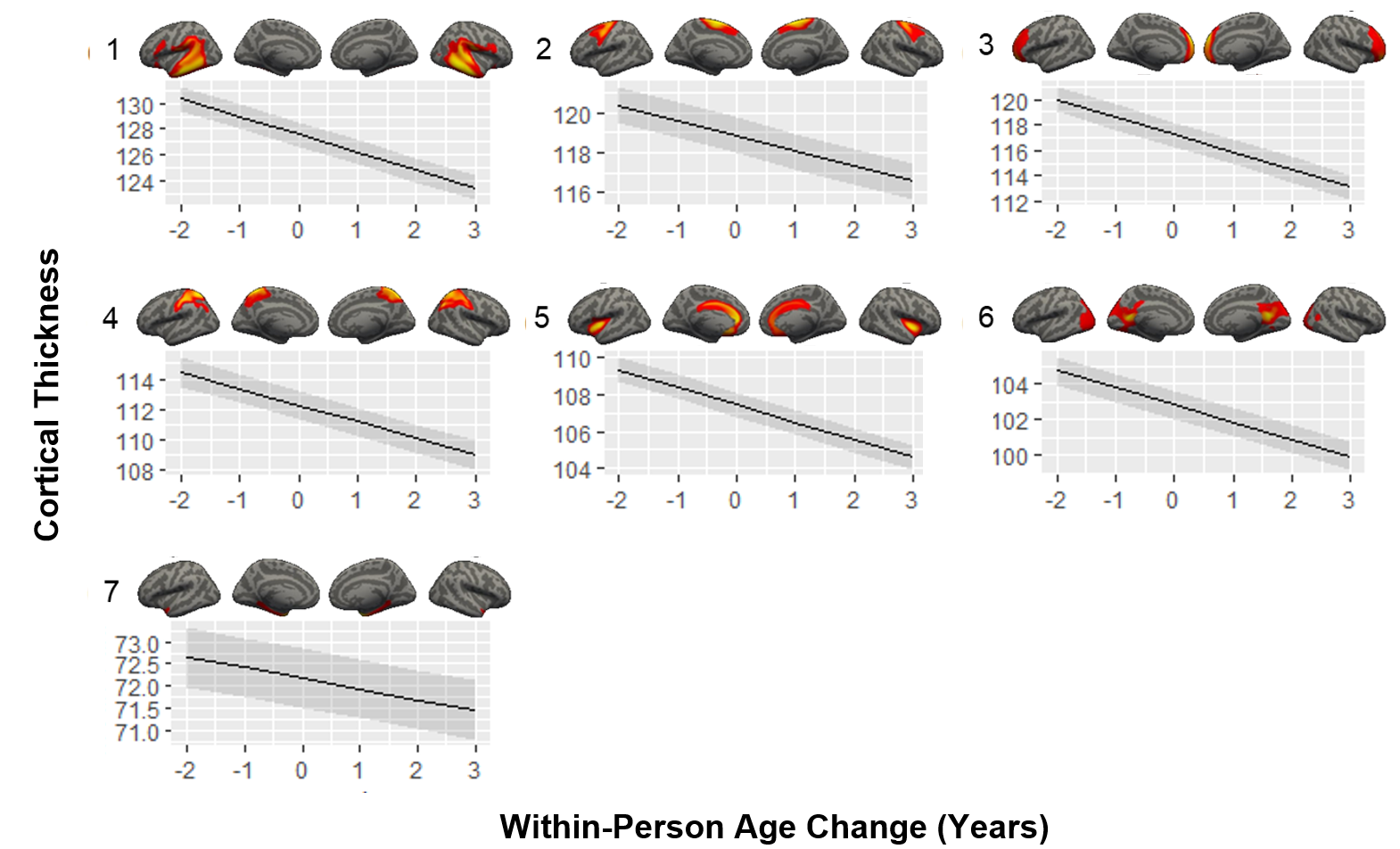
***Supplementary Figure S2.*** Main effect of within-person age change. Older within-person age is associated with lower cortical thickness in all 7 patterns (*β*-values = -1.382~-0.241, t-values = -42.812~-12.166, q-values < 0.001).

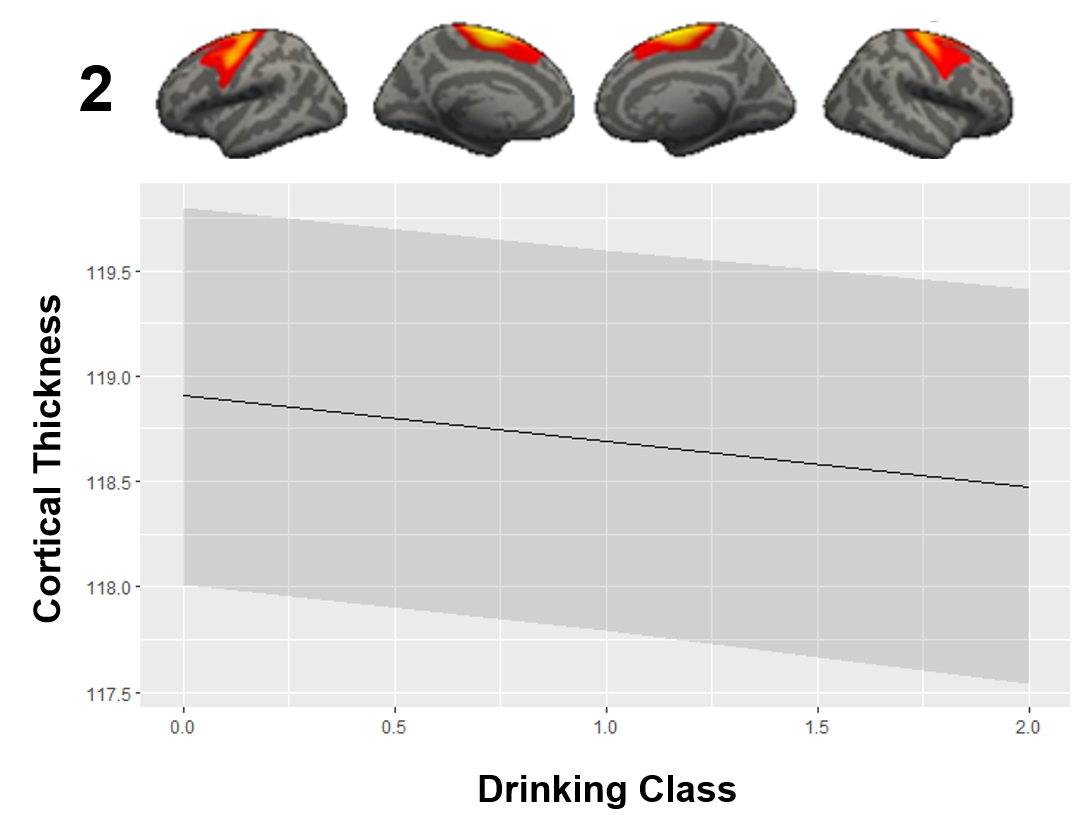
***Supplementary Figure S3.*** Main effect of drinking class. Higher drinking class is associated with lower cortical thickness in pattern 2 (*β*-value = -0.215, *t*-value = -2.752, *q*-value = 0.042).

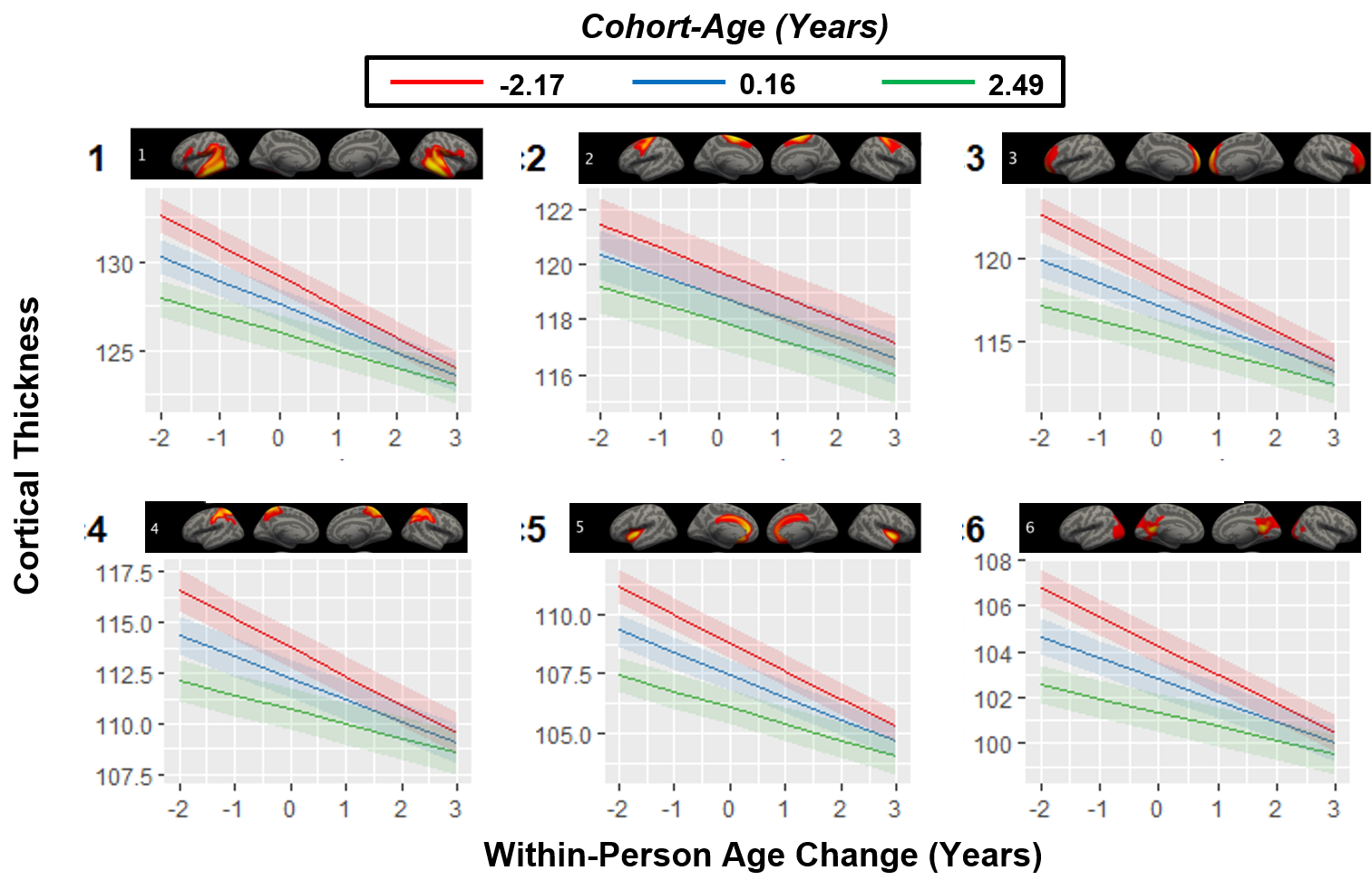
***Supplementary Figure S4.*** Interactions between cohort-age and within-person age change. Participants with older cohort-age relative to those with younger cohort-age are accompanied with slower rate of within-person age change related cortical thickness declines in all patterns (*β*-values = 0.046~0.167, *t*-values = 3.757~13.340, *q*-values < 0.001) except for pattern 7 (*β*-value = 0.012, *t*-value = 1.534, *q*-value = 0.125).

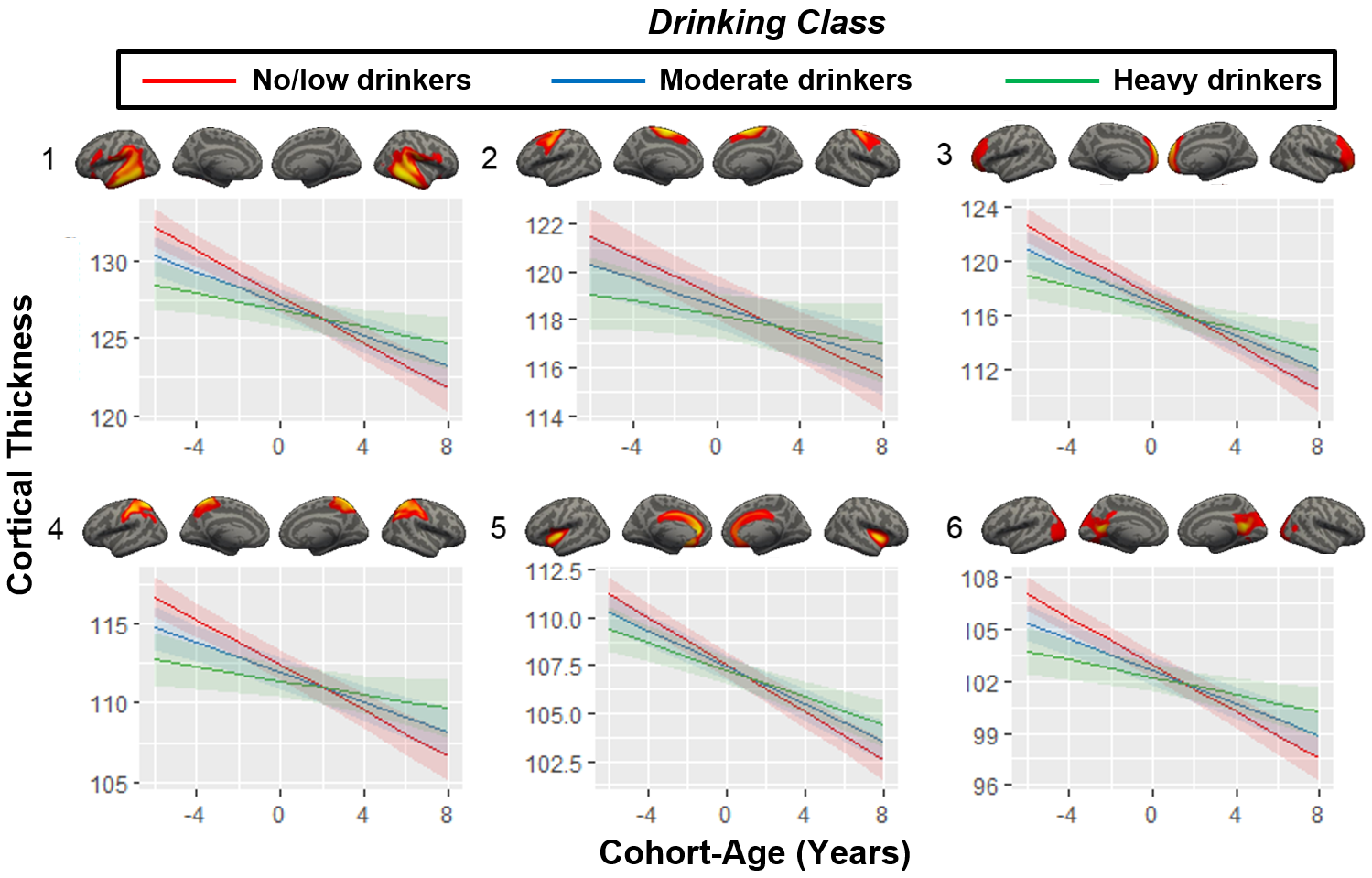
***Supplementary Figure S5.*** Interactions between drinking class and cohort-age. Higher drinking class is associated with slower rate of cohort-age related cortical thickness declines in all patterns (*β*values = 0.134~0.249, *t*-values = 3.887~6.867, *q*-values < 0.001) except for pattern 7 (*β*-value = 0.031, *t*-value = 1.386, *q*-value = 0.166).

***Results of The Solution With 9 NMF Patterns***

In the main text, we reported the findings of the optimal NMF solution that has 7 patterns. As shown in **Fig. 1**, the most prominent peak of split-sample reproducibility that is associated with the optimal solution is close to smaller peak associated with 9 NMF patterns (**Fig. 1S**). We reported below the results related with the 9-patterns solution, which are indeed consistent with the results based on the 7-patterns solution.

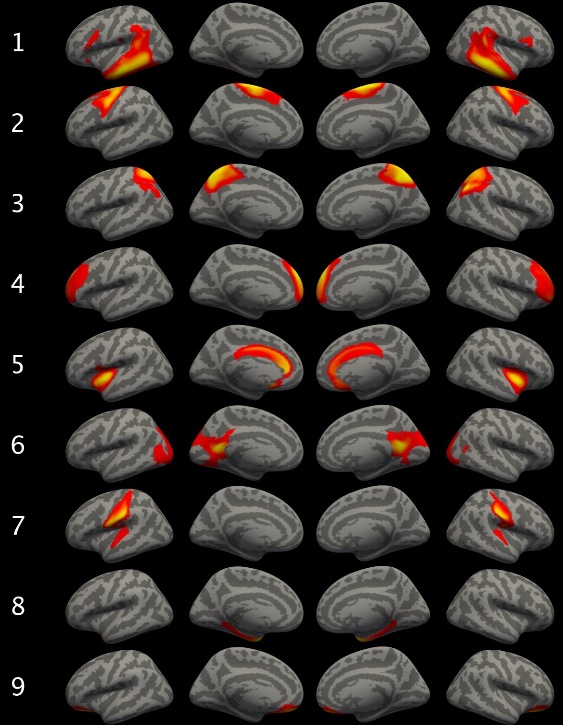

***Figure 1S****. The solution of 9 NMF patterns.*

As shown in **Fig. 1S**, nearly all patterns were highly symmetric bilaterally. Consistent with pattern 1 in the 7-patterns solution, pattern 1 here is mainly associated with voxels in angular gyrus, supramarginal gyrus, inferior frontal areas, and superior/middle/inferior temporal regions. Also consistent with the pattern 2 in the 7-patterns solution, pattern 2 here is mainly related to superior and middle frontal regions. Patter 3 here is similar to pattern 4 in the 7-patterns solution and is associated with superior parietal lobule. Pattern 4 here is similar to pattern 3 in the 7-patterns solution and is associated with frontopolar regions. Pattern 5 is consistent with pattern 5 in the 7-pattern solution and is mainly associated with anterior/middle cingulate cortex and bilateral insula. Pattern 6 is also consistent with pattern 6 in the 7-patterns solution and is associated to posterior cingulate areas and occipital regions. Pattern 7 is related to postcentral regions and superior temporal regions. Pattern 8 here is similar to pattern 7 in the 7-patterns solution and is associated with parahippocampal gyrus. Pattern 9 is associated with the ventromedial prefrontal regions.

*Main Effect of Cohort Age*

Older cohort age was associated with lower cortical thickness in all patterns (*β-values* = -0.746~-0.320, *t*-values = -11.409~-5.652, *q*-values < 0.001) except for pattern 8 (*β-values* = -0.090, *t*-value = -1.821, *q*-value = 0.069).

*Main Effect of Within-Person Age Change*

Older within-person age was associated with lower cortical thickness in all 9 patterns (*β-values* = -1.274~-0.167, *t*-values = -42.815~-8.852, *q*-values < 0.001).

*Interaction of Within-Person Age Change x Cohort Age*

Significant interaction between within-person age change and cohort age was found in all patterns (*β-values* = 0.040~0.161, *t*-values = 3.391~13.141, *q*-values < 0.001) except for pattern 8 (*β-values* = 0.005, *t*-value = 0.727, *q*-value = 0.467). Participants with older cohort age relative to those with younger cohort age are accompanied with slower rate of within-person age-related cortical thickness declines.

*Main Effect of Drinking Class*

No significant effect of drinking class was found in any pattern (*β-values* = -0.162~0.053, *t*-values = -2.701~0.912, *q*-values > 0.06).

*Interaction of Drinking Class x Cohort Age*

Significant interactions were found in all patterns (*β-values* = 0.102~0.261, *t*-values = 3.772~6.821, *q*-values < 0.001) except for pattern 8 (*β-values* = 0.020, *t*-value = 0.957, *q*-value = 0.339). Higher drinking class is associated with slower rate of cohort age-related cortical thickness declines.

*Interaction of Drinking Class x Within-Person Age Change*

No significant result was found in any pattern (*β-values* = -0.064~0.147, *t*-values = -1.458~2.399, *q*-values > 0.1).

*Interaction of Drinking Class x Life Trauma*

No significant result was found in any pattern (*β-values* = -0.065~0.052, *t*-values = -1.599~1.043, *q*-values > 0.8).

*Interaction of Drinking Class x Sex*

No significant result was found in any pattern (*β-values* = -0.260~0.076, *t*-values = -2.092~0.779, *q*-values > 0.3).

*Interaction of Drinking Class x Ethnicity*

No significant result was found in any pattern (*β-values* = -0.402~0.199, *t*-values = -1.360~1.223, *q*-values > 0.7).

*Interaction of Drinking Class x SES*

No significant result was found in any pattern (*β-values* = -0.283~0.136, *t*-values = -1.390~0.465, *q*-values > 0.8).

*Interaction Drinking Class x Family History of AUD Density*

No significant result was found in any pattern (*β-values* = 0.035~0.212, *t*-values = 0.318~1.246, *q*-values > 0.6).

*Interaction of Drinking Class x Cohort Age x Within-Person Age Change*

Significant three-way interactions were found in all patterns (*β-values* = 0.060~0.179, *t*-values = 2.699~7.759, *q*-values < 0.01) except for pattern 8 (*β-values* = 0.005, *t*-value = 0.368, *q*-value = 0.713). The rates of within-person age-related cortical thickness declines are similar across age cohorts in no/low drinkers, faster from the younger cohort and slower from the older cohort in moderate drinkers, and fastest from the younger cohort and slowest from the older cohort in heavy drinkers.
